# Neck motor unit activity displays neural signatures of temporal control during sequential saccade planning

**DOI:** 10.1101/2021.08.31.458347

**Authors:** Debaleena Basu, Naveen Sendhilnathan, Aditya Murthy

## Abstract

Goal-directed behavior involves the transformation of neural movement plans into appropriate muscle activity patterns. Studies involving single saccades have shown that a rapid, direct pathway links saccade planning in frontal eye fields (FEF) to neck muscle activity. It is unknown if the rapid connection between FEF and neck muscle is maintained during sequential saccade planning. We show that sequence planning signals in the FEF are preserved in the neck EMG, although the activity is delayed specifically for the second saccade. Our results suggest that while the direct link between FEF and neck muscle facilitates downstream continuation of FEF response patterns, an indirect route exists through an inhibitory control center like the basal ganglia, limiting the information flow during processing of saccade sequences. Thus, the indirect and direct pathways from the FEF may function together to enable rapid synchronous, but controlled eye-head responses to sequential gaze shifts.

## Introduction

When exploring the visual environment, we move our eyes and head to aim the line of sight (i.e., gaze) at a target of interest for it to be inspected by the eye’s fovea (the area of maximum visual acuity). To achieve accurate gaze shifts, head movements must be precisely coordinated with saccadic eye movements. The frontal eye fields (FEF) have been shown to be a critical neural node for such eye-head coordination (Chen et al., 2006; Constantin et al., 2004; Elsley et al., 2007; Knight and Fuchs, 2007; Martinez-Trujillo et al., 2003; Monteon et al., 2010; Tu and Keating, 2000). However, the link between FEF and neck muscle has not been explored in the context of sequential eye movements - which comprise much of our daily behavior - and forms the basis for this study.

With the head unrestrained, microstimulation of the FEF evokes both eye-only and head-only movements, along with combined eye–head gaze movements, depending on the starting positions of the eyes and the head (Chen et al., 2006; Knight and Fuchs, 2007; Monteon et al., 2010; Tu and Keating, 2000). FEF stimulation results in rapid pre-saccadic recruitment of neck muscle activity, even when there is no overt head movement. Neck muscle recruitment has been shown for subthreshold saccadic stimulation and for small amplitude (∼13°) saccades typically not associated with head-motion (Corneil et al., 2010; Elsley et al., 2007). The activity of the neck muscles, which bring about head movement, is closely linked with saccade generation and the neural activity in FEF (Bizzi et al., 1972b, 1972a, 1971). Finally, the activity of the splenius capitis neck muscle has been shown to be predictive of saccadic RTs in both humans and monkeys (Goonetilleke et al., 2015; Rungta et al., 2021). Together, these studies indicate that FEF neural activity is tightly linked to neck muscle activity by a cascade of rapid downstream events.

Among the various control schemes hypothesized to explain the link between the eye and the head systems, the most parsimonious one suggests that a common gaze displacement signal is separated into eye and head displacement signals before reaching the brainstem movement generator circuitry (**Fig. 1A**). Results from various experimental studies on gaze shifts have converged towards this model (reviewed by Freedman, 2008). Interestingly, selective inhibitory gating of the saccadic burst neurons by the omnipause neurons (‘Gate 2’ in **Fig. 1A**) delays the activation of saccadic burst neurons, while allowing head movement signals to pass through, resulting in the observed presaccadic neck motor unit EMG activity. The early neck EMG signals allow for coordinated eye and head movements during gaze displacement despite the larger inertia of the head, when compared to the eyes. This link persists even during tasks where eye movements were executed in head-restrained conditions, wherein neck EMG activation towards causing a head movement is not required (Lestienne et al., 1984; Rungta et al., 2021). Previous studies evidencing the tight association between neck muscle responses and the FEF have been performed from tasks involving isolated, single saccades. The question of how planning of sequential saccades affects neck muscle activity is yet unknown. Here, we investigated if the link between FEF and neck muscle permitted signatures of sequential saccade planning observed in the FEF, to pass down to the motor periphery.

**Figure 1.**
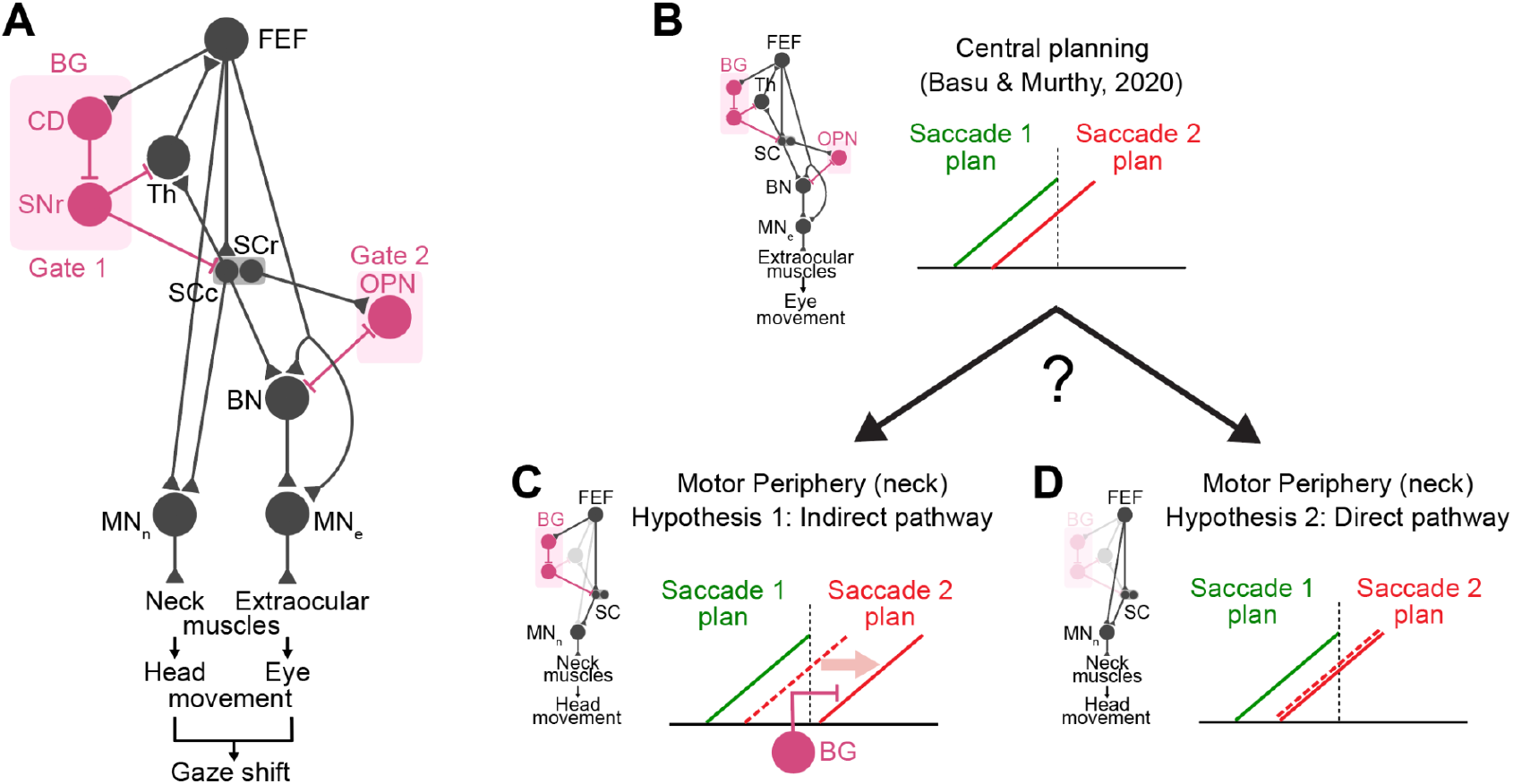
Connections linking FEF to neck muscles. **A**. A basic schematic showing the putative connection between FEF and neck muscle. A common gaze command is relayed from the FEF to the superior colliculus (SC), which is then decomposed into head and eye commands by distinct brain stem premotor circuits. Head premotor cells innervating neck muscles (MN_n_) are not subjected to the inhibition of omnipause neurons (OPN), allowing the rapid pre-saccadic recruitment of neck muscles. BN and MN refer to burst neurons and motoneurons, respectively. The pathway from FEF to superior colliculus is gated by basal ganglia (BG), a major node of inhibitory control. CD and SNr refer to the caudate nucleus and substantia nigra. Connections with triangular endings represent excitatory connections and connections with line endings represent inhibitory connections. **B**. FEF movement activity for the second saccade plan could rise before the onset of the first saccade (as shown in Basu and Murthy, 2020). **C**. Hypothesis 1 - Indirect pathway: If basal ganglia is gating the transmission of motor signals from FEF to neck muscle for sequential saccades, then presaccadic EMG activity related to the second saccade will be delayed or inhibited compared to the FEF signal. A possible anatomical framework explaining this is shown to the left. **D**. Hypothesis 2 - Direct pathway: If the basal ganglia gate does not play a major role, then a direct connection between FEF and motor neck muscles (as shown in the left) could allow for EMG activity to be initiated with minimal delay after the onset of presaccadic FEF activity. The EMG activity would be expected to mirror the FEF activity patterns.

Neck muscle activity during single saccade planning has been shown to follow FEF activity closely with a short latency (Elsley et al., 2007). We have previously shown that a sequence of two saccades may be programmed in parallel in the FEF, albeit with processing limitations (**Fig. 1B**; Basu et al., 2021; Basu and Murthy, 2020). The transfer of concurrent saccade programming signals from FEF to neck muscle may be inhibited to prevent premature head motion for second gaze shift while the first gaze plan is still underway. The node for such inhibitory control may be at the basal ganglia, which has been shown to bring about processing bottlenecks to prevent potential detrimental consequences of parallel programming of saccade plans (Bhutani et al., 2013). Basal ganglia also sends inhibitory projections to the superior colliculus (Hikosaka and Wurtz, 1985a, 1985b, 1983) directly through the substantia nigra pars reticulata (SNr) nuclei. Given its role in inhibitory control (Aron et al., 2007; Brittain et al., 2012; Frank et al., 2007), and sequential motor control (Aldridge and Berridge, 1998; Kermadi and Joseph, 1995; Mushiake and Strick, 1995), basal ganglia might act as a possible inhibitory control node for gaze shifts, delaying or inhibiting the downstream leakage of signals associated with the second saccade plan when programmed in parallel (*hypothesis 1* **Fig. 1C**). Alternatively, in a scenario mimicking the planning of single saccades, FEF signals encoding saccade sequences may pass down freely to the neck musculature with minimal conductive delay and bring about pre-saccadic recruitment of neck motor units (*hypothesis 2* **Fig. 1D**).

Our results show that aspects of both the direct and indirect connections between the FEF and neck muscle come into play during sequential saccade planning. Even when head movements were not required or executed, neck motor unit EMG showed remarkably preserved presaccadic activity profiles: signatures of parallel programming and its consequent processing bottlenecks in the FEF were observed in the peripheral neck EMG. The congruence between neural and peripheral activity profiles supports the hypothesis that a direct channel exists between FEF and neck muscle, even for planning of saccade sequences. However, the onsets of EMG activity were significantly delayed compared to neural activity onsets especially for the second saccade plan, indicating that downstream flow of signals through the FEF-neck EMG circuit is not free from inhibitory control.

## Results

Two monkeys performed a sequential saccade task (FOLLOW task; see Methods) where in 70% of the trials (called ‘step trials’; **Fig. 2A**), they performed a rapid sequence of visually-guided saccades to two targets (T1 and T2) in the order of their appearance. In the step trials, the time between the T1 and T2 (called the target step delay or TSD) was randomly chosen among 17 ms, 83 ms, and 150 ms on each trial. In the remaining 30% of the trials (called ‘no-step’ trials; **Fig. 2B**), they made a single saccade to a single visual target that was presented. The two types of trials were randomly interleaved.

**Figure 2.**
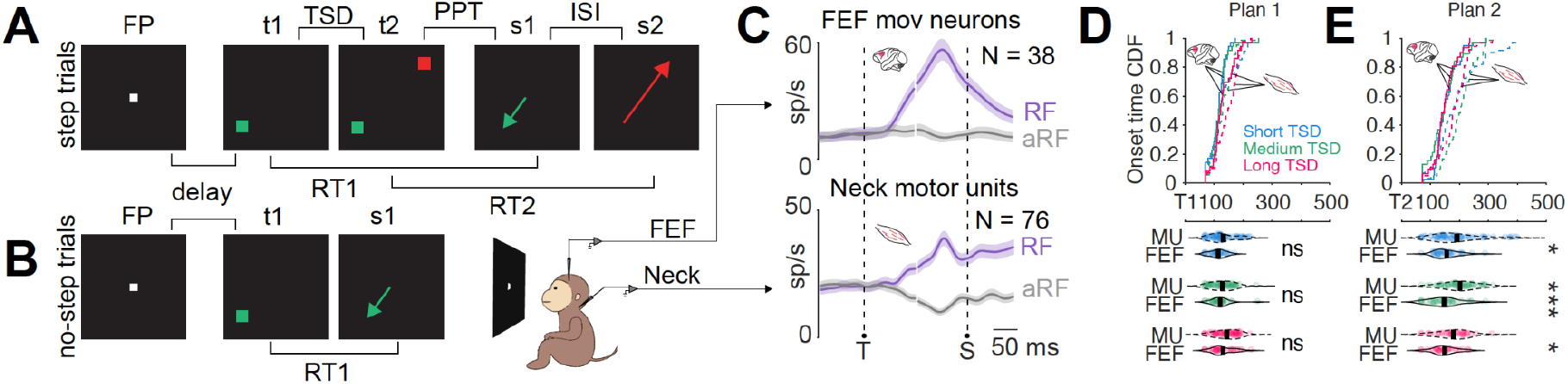
FEF and EMG Activity in the FOLLOW task. **A**. Schematic of a step trial in the FOLLOW task. **B**. Schematic of a no-step trial in the FOLLOW task. **C**. Population activity for saccades into the response field (RF; *purple*) and saccades out of the response field (aRF; *gray*) for FEF movement neurons (top) and neck motor units (bottom) aligned to target onset (T) and saccade onset (S) during the no-step trials. The solid line indicates the mean activity, and the shaded region indicates the mean ± SEM. **D**. *Top*: CDF of onset times for three target step delays (TSD) for FEF and motor units for saccade plan 1. *Bottom*: Same data as top panel visualized in bee-swarm plots. The vertical black line within each plot indicates the average value. **E**. Same as **D** but for saccade plan 2.

As the monkeys were performing the task, we recorded from FEF neurons and motor units from the dorsal neck muscle. We selected 76 motor units that were well isolated for further analyses. During a memory-guided saccade task (see Methods; **Fig. S1)**, all the motor units showed a significant pre-saccadic rise of activity when the saccades were made into their response field (see Methods; **Fig. S1**) compared to when the saccades were made opposite to their response field. However, they had little or no activity in the visual epoch regardless of whether the target was present in the response field or not. Similar activity patterns were seen in no-step trials as well (**Fig. 2C)**. To interpret the results in the context of FEF responses, we analyzed FEF neurons with a similar activity profile, i.e., FEF movement neurons (**Fig. 2C)**.

Previous experiments have shown that microstimulation of the FEF elicits short latency (∼20 ms) EMG responses from the dorsal neck muscles during a single saccade task (Elsley et al., 2007). To check the latency of neck muscle responses during sequential saccade planning, we compared the onsets of presaccadic activity for FEF and neck EMG for each of the two saccades in the FOLLOW task. In the trials where the first saccade went into the response field, the onsets for FEF and EMG activity were not significantly different at the population level for each of the three target step delays (*short*: Kruskal-Wallis, *χ*^*2*^ (1, 101) = 2.58, *p* = 0.11; *medium*: Kruskal-Wallis, *χ*^*2*^ (1, 97) = 1.4, *p* = 0.24; *long*: Kruskal-Wallis, *χ*^*2*^ (1, 92) = 3.46, *p* = 0.06). Activity for the second saccade plan on the other hand showed significantly different onsets between EMG and FEF (*short*: Kruskal-Wallis, *χ*^*2*^ (1, 108) = 4.77, *p* < 0.05; *medium*: Kruskal-Wallis, *χ*^*2*^ (1, 96) = 20.15, *p* < 0.001; *long*: Kruskal-Wallis, *χ*^*2*^ (1, 66) = 6.82, *p* < 0.01). **Fig. 2D-E** shows the cumulative distribution function of the activity onset times for neck EMG and FEF. While comparing onsets is an indirect measure of the delay between FEF and neck EMG, our results show that the differences in activity onsets is amplified specifically for the second saccade plan, indicating that inhibitory control comes into play in the FEF-neck EMG circuit for multiple saccades planned in a sequence.

### Peripheral signatures of parallel programming during sequential saccades

Even though presaccadic neck EMG activity showed delayed onsets for the second saccade plan and implicates the indirect circuit (**Fig. 1C)**, neural signatures of sequential saccade planning may still pass down from the FEF to the motor periphery, albeit with a delay. Our previous study has shown neural correlates of parallel programming of saccades in the FEF (Basu and Murthy, 2020). To assess whether neck motor activity carries signatures of parallel programming, we first analyzed saccadic behavior during the EMG sessions. The nature of sequential planning can be assessed by testing whether the interval between the saccades varies systematically with the duration available for parallel programming. This duration, called the parallel processing time (PPT; similar to delay D of Becker and Jürgens, 1979), is the time when both saccade plans are underway, i.e., the time period from the appearance of the second target, to the end of the first saccade. If saccade programming were strictly serial, the inter-saccadic interval (ISI) would be fixed and independent of the time available for parallel programming (long or short PPT; **Fig. 3B**). In contrast, if the second saccade can be planned in parallel, shorter ISIs can be obtained for larger PPTs (**Fig. 3B**). Thus, the slope of the ISI-PPT plot is a behavioral metric for the concurrent programming of saccades (Becker and Jürgens, 1979; Bhutani et al., 2013, 2012; McPeek et al., 2003, 2000; Minken et al., 1993; Ray et al., 2004; Sharika et al., 2008; Wu et al., 2013). Consistent with this notion, the inter-saccadic interval (ISI) decreased significantly as the parallel processing time increased (**Fig. 3C**; one-way ANOVA, F (1, 148) = 189.82, *p* < 0.001). Each session showed slopes that were significantly below zero (**Fig. 3C** inset; two-sided Wilcoxon signed rank test, Z = 5.21, *p* < 0.001).

**Figure 3.**
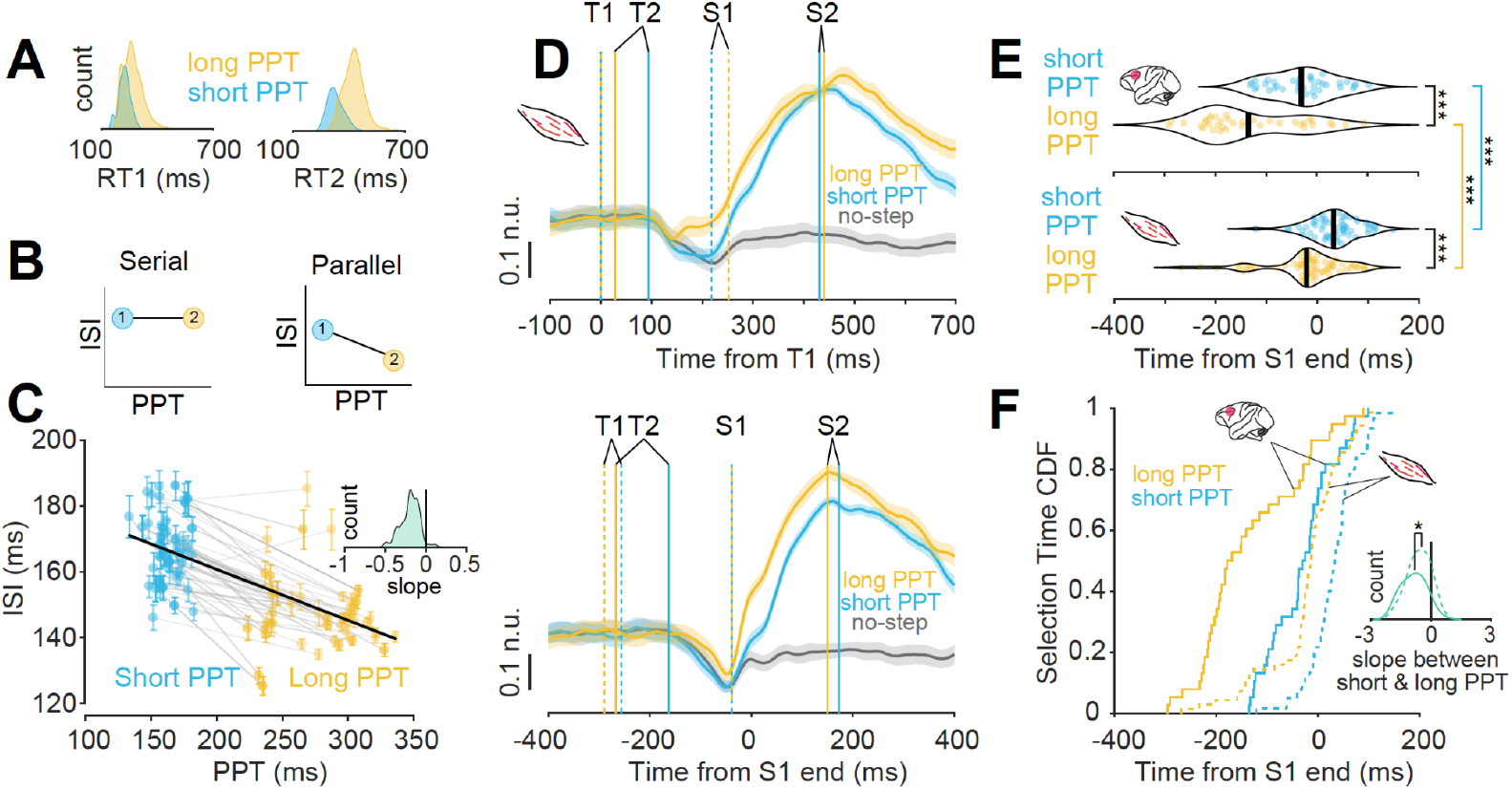
Activity of neck muscle motor units show correlates of parallel programming. **A**. RT1 distribution (top) and RT2 distribution (bottom) for short (blue) and long (yellow) PPT conditions. **B**. Illustration of the relationship between ISI and PPT for serial (top) and parallel (bottom) processing. **C**. ISI vs PPT plot for each session. Inset shows the distribution of slopes between ISI and PPT. **D**. Top: Motor unit population activity (mean ± SEM) when the second saccade was made into the movement field for short and long PPT conditions (*blue* and *yellow* respectively). The activity is aligned to the first target onset and contrasted with the no-step activity of saccades made outside of the movement field (*black line*). T1 and T2 represent target 1 and target 2 onsets and S1 and S2 represent saccade 1 and saccade 2 onsets. Bottom: same as top panel but aligned to the end of the first saccade. The solid line indicates the mean activity, and the shading indicates mean ± SEM. **E**. Top: Neural selection times for all the FEF neurons for short and long PPT (as already shown in Basu and Murthy, 2020). Bottom: Peripheral selection times for all the motor units for short and long PPT. *** means *p*<.001 **F**. Same data from **E**, represented as cumulative selection times for short (*blue*) and long (*yellow*) PPTs for motor units (*broken lines*) and FEF neurons (*solid lines*). Inset: The distribution of slopes between short and long PPT for FEF neurons (*solid line*) and motor units (*broken line*). * means *p*<0.05

We then evaluated if the neck EMG activity showed correlates of parallel programming by analyzing the trials in which the second saccade went into the response field. Parallel programming at the level of the responses of FEF movement neurons was evinced by two main activity trends (Basu and Murthy, 2020): First, the neural selection time, demarcating the onset of neural activity specific to the second saccade, could start before the visual feedback of the new eye position after the execution of the first saccade could reach FEF. Often the neural activity related to the second saccade started much before the onset of the first saccade, showing clear evidence that movement activity of a saccade plan can ramp-up while a previous plan is still ongoing. Second, as the parallel programming time (PPT) available decreased, the neural selection times progressively got delayed with respect to the first saccade onset, thus providing a direct correlation between neural activity and behavioral markers of parallel programming (Basu and Murthy, 2020).

If the basal ganglia circuitry restricts the signal flow between FEF and neck muscle for the second saccade, then it is plausible that activity at the level of neck muscle will be serialized (**Fig. 1C**). Neck muscle activity, however, showed remarkably preserved signatures of neural correlates of parallel programming, despite head-restraint and no requirement for overt head movements. The peripheral selection time (PST), calculated analogously to the neural selection time (NST; Basu and Murthy, 2020), was estimated by calculating the time when EMG activity for the second saccade plan crossed two standard deviations above the baseline activity (see Methods). The trials in each session were divided into short and long PPT trials based on whether the PPT value in a trial was below or above the mean PPT of the session. The peripheral selection times occurred before the visual feedback latency and sometimes even before the onset of the first saccade, especially for trials in the long PPT condition), indicating that motor unit activity associated with the second saccade plan could emerge before the end of the first saccade (**Fig. 3D**; median PST in long PPT group = -12 ms). Further, the peripheral selection times in the long PPT group were significantly lower than those of the short PPT group, i.e., when the time available for parallel programming is higher, the peripheral selection time shifts earlier in time with respect to the first saccade (**Fig. 3E**, Kruskal-Wallis, χ^2^ (1, 127) = 21.92, *p* <.001) similar to the pattern seen in the concurrent FEF neural activity (Basu and Murthy, 2020). At the population level, the average peripheral selection time slope was significantly less than zero (two-sided Wilcoxon signed-rank test, *Z* = -5.66, *p* <.001), thus corroborating with the inverse NST-PPT relation shown in our previous study. Similar peripheral correlates of parallel processing were obtained for a restricted set of sessions with reaction time matched trials (**Fig. S2**), indicating that the inverse PST-PPT relationship we observed is not just a result of stochastic variability in the saccade planning processes. Thus, EMG activities elicited by multiple saccade plans can be active simultaneously, similar to the FEF activity patterns.

However, the peripheral selection time for motor units and neural selection time for FEF neurons differed significantly in the short (Wilcoxon signed-rank test, *p* < 0.001) as well as long (Wilcoxon signed-rank test, *p* < 0.001) PPT conditions (**Fig. 3E**). The NST-PPT slopes were also more negative (−0.85 ± 0.01) than the PST-PPT slopes (−0.59 ± 0.01; Two-sample t-test *p* < 0.05) suggesting that the EMG activity was less parallel than the FEF. The pattern of the cumulative distributions of the neural selection times of FEF movement neurons and peripheral selection times of neck muscle motor units shows that the peripheral selection times were delayed compared to the neural selection times (**Fig. 3F**). The median neural selection time in FEF for the long and short PPTs were 161 and 62 ms earlier compared to the peripheral selection times.

Thus, neck EMG activity can ramp up for a consecutive saccade whilst the previous one was still being programmed, fitting into hypothesis 2 (**Fig. 1D**), which is consistent with direct feedforward activation from FEF to the neck motor units. Paradoxically, EMG activity for the second saccade plan was delayed substantially with respect to the FEF activity onsets, providing evidence for hypothesis 1 (**Fig. 1C**). Our results indicate that while signals encoding saccade sequences pass down from oculomotor centers to the motor periphery and show similar patterns, the flow is not unchecked: the second saccade plan especially, appears to be gated by inhibitory control centers and thus has delayed activity onsets.

### Peripheral signatures of processing bottlenecks during sequential saccades

A complimentary aspect of parallel planning that enables rapid saccade sequencing, is the idea of processing bottlenecks that limit the extent of parallel planning so that saccades that are planned together are prevented from being executed together resulting in errors like averaged saccades, altered saccade metrics, or incorrect saccade order (Bhutani et al., 2017, 2012). An increase in movement latencies provides behavioral evidence of processing bottlenecks (Pashler, 1994). Consistent with this notion, for both the first and second saccades, the reaction times across the population increased as the target step delay decreased from 150 to 17 ms (RT2: one-way ANOVA, F (2, 160) = 112.09, *p* < .001; **Fig. S2**). In our previous study, we have shown that these longer response times reflect the activity of FEF responses which slows down when two closely-spaced saccade plans proceed simultaneously (Basu et al., 2021).

Before checking EMG responses for sequential saccades, we checked the EMG responses for single saccades in the no-step trials. Corroborating with the rise-to-threshold hypothesis of accumulator models (Hanes and Schall, 1996), FEF movement neuron activity altered the rate of growth but not the threshold to account for reaction time variability in the single saccade no-step trials (Basu et al., 2021). The EMG activity showed similar profiles, ramping up activity in an accumulator framework, with the rate of growth on an average, being greater in trials with faster reaction times (one-way ANOVA, F (1, 123) = 12.89, *p* < .001). The threshold activity did not change among the slow and fast reaction time groups (**Fig. S4**).

Having verified that the pattern of single-saccade related activity was preserved from FEF to the neck muscle, we next looked at the activity of motor units during step trials during sequential saccades. The accumulator framework for explaining reaction time variability observed in saccadic tasks have mostly been used for single saccade tasks. Our previous study showed that for sequential saccades, the accumulator parameters of rate and threshold varied with target step delay, but onset and baseline activity did not. We performed a similar analysis for neck EMG data to check if central planning signals reaching the motor periphery followed the pattern of accumulator activity observed in FEF.

EMG activity in trials in which the second saccade went into the RF distinguished between the target step delay conditions in a manner similar to FEF movement neurons: the rate of neck muscle activity decreased with decrease in target step delay, whereas threshold increased (**Fig.4A**). We performed a regression analysis for each motor unit wherein the slope of the best fit line was taken for each parameter (see Methods). Only the slopes for rate of activity growth and threshold for motor units were significantly different from zero (Wilcoxon signed-rank test for slopes of rates, Z = -2.71, *p* < .01, **Fig. 4B**; Wilcoxon signed-rank test for threshold of threshold, Z = -3.64, *p* < .01, **Fig. 4B**. Baseline and onset of activity did not distinguish between the target step delays across the population (*p* > .05). The changes in these parameters reflected the population dynamics seen at the FEF level (Basu et al., 2021; **Fig. 4C**). However, the slopes for rates were significantly higher for FEF neurons compared to motor units (Two-sample t-test *p* < .05). The slopes for thresholds were not significantly different between FEF neurons and motor units (Wilcoxon signed-rank test *p* = 0.3368).

**Figure 4.**
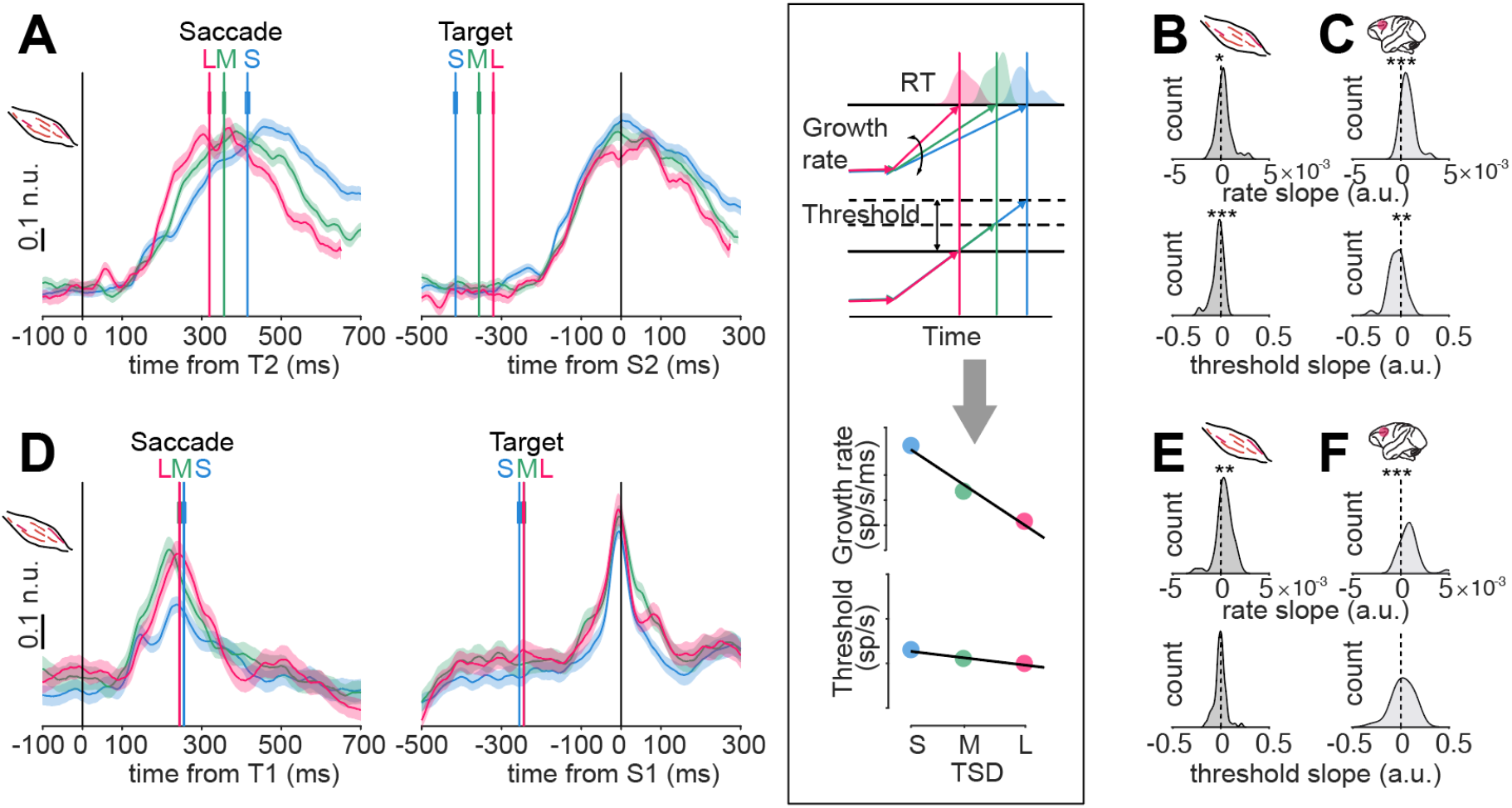
Activity of motor units during the second saccade plan show signatures of processing bottlenecks. **A**. Population activity of the motor units when the second saccade went into the response field, aligned on the second target onset (*left*) and the start of the second saccade (*right*). The solid line indicates the mean activity, and the shaded area indicates mean ± SEM. S, M, L indicate short, medium, and long TSD respectively. *Boxed inset*: The top panel shows the modulations expected for growth rate and threshold activity, along with the RT histograms. The bottom panel shows a schematic of the variation in growth rate and threshold across TSD for an example motor unit. For each motor unit, the slope derived for each parameter from the best fitting line is used in Figures B & E. **B**. Histograms of slopes of the change of muscle activity parameters (*top*: rate, *bottom*: threshold) with target step delay in individual motor units. **C**. Same as **C** but for FEF neurons as already shown in Basu et al., 2021. **D.** Population activity of the motor units when the first saccade went into the response field, aligned on the first target onset (*left*) and the start of the first saccade (*right*). The solid line indicates the mean activity, and the shaded area indicates mean ± SEM. **E.** Histograms of slopes of the change of muscle activity parameters (*top*: rate, *bottom*: threshold) with target step delay in individual motor units. **F**. Same as **F** but for FEF neurons as already shown in Basu et al., 2021.

Consistent with the results obtained from the FEF movement neuron activity (Basu et al., 2021), rate perturbation was present even when the first saccade went into the response field (**Fig. 4D-E**; Wilcoxon signed-rank test, Z = -3.54, *p* < .0001). The slope of the threshold was not significantly different from zero (t-test p = 0.3090; **Fig. 4E**) similar to FEF neural activity (t-test *p* = 0.2910; **Fig. 4F**). However, the slopes for rates were slightly higher for FEF neurons compared to motor units (Two-sample t-test *p* = 0.0446). The slopes for thresholds were not significantly different between FEF neurons and motor units (Wilcoxon signed-rank test, *p* = 0.1777). The fact that rate perturbation was present in both the saccade plans indicates that the signatures of processing bottlenecks that were observed in the responses of FEF neurons, were also seen in the motor periphery consistent with hypothesis 1 (**Fig. 1C**). Thus, neck muscle EMG patterns closely followed that of FEF, even though neck EMG activation for any oncoming head movement was unnecessary due to head-restraints.

## Discussion

In this study, we investigated the recruitment of the dorsal neck muscle in monkeys making sequential visually-guided saccades. The results indicate that putative motor units encoding the anticipated gaze movement for the second movement in a sequence are recruited in parallel with those encoding the first gaze shift vector, just as it is observed in FEF movement neurons. However, inhibitory control, specific for the second movement, was more prominent in the activity of motor units compared to the FEF. Taken together, these results suggest signatures of both the direct and indirect pathways between the FEF can be observed in the activity of neck muscles. The tight downstream linking of the FEF and the motor periphery is emphasized by the fact that neural patterns are preserved in the periphery even when no peripheral EMG activation is required by the task.

### Peripheral correlates of gaze planning signals

The direct linking hypothesis between the FEF and neck musculature (Corneil et al., 2010, 2002a, 2002b) assumes that the FEF encodes a composite gaze command, which is then relayed to the superior colliculus, following which the eye and head commands are decomposed and relayed to separate brainstem premotor circuits. This scheme is supported by the fact that electrical stimulation of the FEF and caudal superior colliculus produces eye and head movements in cats and monkeys (Chen, 2006; Corneil et al., 2010, 2010; Cowie and Robinson, 1994; Elsley et al., 2007; Freedman et al., 1996; Harris, 1980; Isa and Sasaki, 2002; Roucoux et al., 1980; Stryker and Schiller, 1975; Tu and Keating, 2000). Downstream to the superior colliculus, these gaze-related motor commands drive the extraocular and neck muscles, respectively, producing a coordinated gaze shift. Such coordination is thought to be facilitated by the absence of inhibitory gating by the omnipause neurons (Gate 2 in **Fig. 1A**) which may allow activity development in the inertia-laden neck muscle, leading to its rapid pre-saccadic recruitment, while preventing the signal from reaching the eye muscles prematurely. This scheme forms the basis of our observation of a central correlate of movement planning reported in this study.

One prediction of such a direct pathway is that the latency of evoked neck muscle responses following FEF microstimulation can be as low as 20 ms (Elsley et al., 2007), probably indicating a ‘cephalomotor expression of a transient visual response that sweeps through extrastriate and oculomotor areas shortly after visual target onset’ (Goonetilleke et al., 2015). In this context, Corneil et al., (2004), found short latency stimulus-locked muscle responses, while the visuo-motor index of the motor units collected for this study (see Methods) was close to being pure movement -- in other words, the activity was saccade-related and did not show the visual burst. One explanation for the lack of visual activity is that it may be triggered in the context of a grasp reflex that demands rapid orienting of gaze (Corneil et al., 2008, 2004). As noted before, such leakage of signals into the periphery may also be sensitive to context (Pruszynski et al., 2010; Wood et al., 2015). Another and possibly more likely explanation could be that even in the original study done by Corneil et al. (2004), the stimulus-locked response of the splenius capitis muscle was much lower, while deeper muscles such as the rectus capitis posterior and the obliquus capitis inferior muscles showed robust stimulus-locked responses. Our recordings were done using external landmarks on the dorsal neck plane (see Methods) and we targeted the splenius capitis muscle, a large, superficial, ipsilateral head-turner, as it is easily accessible from the surface. When targeting the splenius capitis, the possibility of penetrating the rectus capitis posterior and obliquus capitis inferior is present but is highly unlikely as these smaller, deep-set muscles are difficult to reach. Nonetheless, despite the absence of a robust visual response, these results show that motor unit responses related to the second saccade can get initiated before the first saccade is completed.

An additional difference between current work and previous work is that the EMG signals were processed using a raster-based method, which is the standard approach used for neural data to answer how information is represented in different brain areas. We used the same approach to compare activity between the FEF and the neck motor unit activity in the periphery. In this context, it is interesting to note that a simple accumulator framework appeared to fulfil the requirements of a unifying framework that could link central processes like movement preparation to recruitment of motor units from periphery and behavioral reaction times **(Fig. S4**; Basu and Murthy, 2020; Carpenter and Williams, 1995; Hanes and Schall, 1996; Ramakrishnan et al., 2010). In such a framework, the rate of accumulation to a constant threshold determines the reaction time and forms the basis of studying their modulation in FEF and the peripheral musculature during sequential movements. Further, the congruence of patterns in FEF and neck muscle activity in relation to reaction time reinforce the claim that such patterns of activity in the periphery reflect central processing as a consequence of the direct pathway between the FEF and the neck musculature.

### Peripheral correlates of inhibitory control of sequential movements through the indirect pathway

While the predictions of the direct circuit are well-tuned to the existing results from single gaze shifts, when extending this circuit from single to sequential movements, inhibitory control pathways specific to controlling sequences might come into play as a part of volitional gaze control. Bhutani et al. (2013) showed that the basal ganglia, a critical node of inhibitory control, is involved in the conversion of parallel movement plans into sequential behavior. Inactivation of the basal ganglia in monkeys or impairment of the basal ganglia in patients of Parkinson’s disease resulted in a significantly greater extent of saccadic errors that develop due to unchecked parallel programming leading to a ‘collision’ of movement plans. It is well established that the connection between the FEF and the superior colliculus, a major sub-cortical node of oculomotor planning, share connections through the output nuclei of basal ganglia (substantia nigra pars reticulata; Hikosaka et al., 2000; Gate 1 in **Fig. 1A**), and possibly, it is this loss of inhibition (transiently by muscimol inactivation or chronically in Parkinson’s disease patients) that led to an increase in saccadic errors. Given the importance of the role of the basal ganglia in the correct execution of saccade sequences, it stands to reason that such inhibitory control specific to sequences of gaze to prevent concurrent activations of the sluggish neck muscles (agonist and antagonist), and any detrimental synergies that might develop from multiple co-activations.

Since the FEF-neck muscle ‘neural highway’ involves the FEF-superior colliculus circuit, which is known to be gated by the basal ganglia (Gate 1 in **Fig. 1A**), the downstream leakage of gaze planning signals from FEF to the motor periphery might be limited by the basal ganglia inhibitory node for more than one gaze shift. Consistent with this hypothesis, our results show neck muscle activity for the second saccade was delayed relative to the neural selection time in FEF for all target step delays but was not significantly different for the first saccade (compare **Figs. 2D** and **2E**).

The selective delaying of the peripheral selection time (PST) for the second saccade results in the shortening of the intervals between PST1 and PST2, relative to NST1 and NST2 (compare **Fig. 4E** and **Fig. 4F)**. In other words, the effective processing rate is smaller in the periphery compared to the FEF. Thus, it appears that the descending input from FEF to superior colliculus, the putative pathway through which the gaze command leaks down to the motor periphery, is routed through the basal ganglia gate specifically to delay and serialize neck muscle responses. However, the inhibition is not complete or strong enough to act as a ‘global stop’ preventing all passive leakage from the FEF or inhibit concurrent flow of gaze planning signals to the periphery, as seen in the results of our study.

### Peripheral signatures of parallel processing and processing bottlenecks for sequential gaze shifts

Although neck muscles responses did show the presence of additional inhibitory control, presumably mediated by the intervening basal ganglia, a rather surprising result was the close correspondence between signatures of planning in the FEF and the motor unit activity. Like in FEF, we found evidence of parallel processing such that motor unit responses related to the second saccade can get initiated before the first saccade is completed (**Fig. 3**). The onsets of the EMG activity, or the peripheral selection times occurred earlier as PPT was increased, similar to the results obtained with neural selection times (Basu and Murthy, 2020; **Fig. 3**). Further when two saccade plans overlap too closely (target step delay = 17 ms), the extent of parallel programming is controlled by adjustments in the rate and threshold of EMG activity in a manner similar to what is seen in FEF movement neurons (**Fig. 4**). These adjustments closely match those of FEF movement neurons: while changes in both slope and rate in FEF and motor unit activity were associated with the second saccade, only changes in slope but not threshold were observed in FEF and motor unit activity were associated with the first saccade response. These results extend the link between the FEF and neck muscle seen in single saccade tasks to sequential saccade planning. The fact that such central planning patterns for sequential saccades are maintained at the periphery suggests that the leakage down of signals at the periphery is almost like a passive down-flow of central planning signals involving the direct pathway. Therefore, taken together, our results suggest that motor unit activity reflects input from both the direct and indirect pathways: while the direct pathway mirrors the activity of the FEF onto the motor units; inhibitory control by the basal ganglia, through the basal ganglia-thalamo-cortical loop, may be another pathway through which the basal ganglia can modulate the activity of FEF neurons and bring about the observed processing bottlenecks in both the FEF and the motor unit activity.

## Acknowledgements

We thank S. Sengupta for helping with behavioral training and Dr. A. Gopal P.A. and Dr. S. Rungta for helping with the data collection.

## Author contributions

Conceptualization, A.M.; Methodology, D.B.; Formal analysis, D.B.; Investigation, D.B.; Writing – Original Draft, D.B., N.S., & A.M.; Writing – Review & Editing, D.B., N.S., & A.M.; Visualization, D.B. & N.S.; Supervision, A.M.; Project administration, A.M.; Funding Acquisition, A.M.

## Methods

The methods that were used in this study have been described in detail elsewhere (Basu et al., 2021; Basu and Murthy, 2020; Rungta et al., 2021; Sendhilnathan et al., 2021). Here, we describe them briefly.

### Subjects

We used two adult monkeys, J (*Macaca Mulata*, male, age = 9 yrs; weight = 5.5 kgs) and G (*Macaca radiata*, female, age = 11 yrs; weight = 3.8 kgs) for the experiments. All surgical procedures and monkey care were in compliance with the animal ethics guidelines of the Committee for the Purpose of Control and Supervision of Experiments on Animals (CPCSEA), Government of India, and the Institutional Animal Ethics Committee (IAEC) of the Indian Institute of Science that approved the protocols.

### Behavioral tasks

The monkeys were trained on two oculomotor tasks: the memory-guided saccade task and the FOLLOW task. In the memory-guided saccade task, each trial started with a red fixation point (0.6° × 0.6°) appearing at the center of a screen. After a variable fixation period, a gray target stimulus (1° × 1°) was flashed briefly (100 ms) at a peripheral location. The monkeys continued fixating for about 1000 ms (delay period), following which the central fixation spot disappeared. A single saccade had to be made to the remembered location of the target, after which juice rewards were given. The MG task was used to identify the response field of neurons (**Fig. S1;** see next section below) and to classify the neurons. In the FOLLOW task (**Fig. 2A, B**) monkeys made a sequence of two visually-guided saccades. After central fixation, a green target appeared at any one of the six possible peripheral locations. In 70% of the trials (*step trials*) the first green target was followed by a second red target and the monkey had to execute a sequence of two saccades in order of target appearance. The remaining *no-step* trials had only one target and the monkey had to make one saccade to the target. The temporal gap between the first and second targets in step trials is referred to as the target step delay and was picked randomly from 17 ms, 83 ms, and 150 ms.

### Data acquisition

The tasks were controlled and displayed using a TEMPO/VIDEOSYNC system (Reflecting Computing, St. Louis, MO, USA). Electrophysiological data was acquired using the Cerebus data acquisition system (Blackrock Microsystems, Salt Lake City, UT, USA). A monocular infrared pupil tracker (ISCAN, Woburn, MA USA) was used to collect eye position data. All stimuli were presented on a Sony Bravia LCD monitor (42 inches, 60 Hz refresh rate; 640 × 480 resolution) placed 57 cm from the monkeys. The monkeys were head-restrained during the tasks.

The electrophysiological data consisted of neural data from the FEF and Electromyographic (EMG) data from the dorsal neck muscles. Neural data was recorded using tungsten microelectrodes (FHC, Bowdoin, ME, USA; impedance: 2 to 4 MΩ). EMG activity from the dorsal neck muscle was recorded bilaterally using intramuscular, Polytetrafluoroethylene-coated stainless steel needle electrodes (diameter 0.36 mm; TECA Elite series, Natus Neurology, Middleton, WI, USA). EMG needle electrodes were inserted using externally available landmarks on the dorsal neck. The dorsal neck plane was framed into a two-dimensional Cartesian coordinate system using the external occipital protuberance and the dorsal midline as the horizontal and vertical axes respectively. All the insertions for the two monkeys were within 2-4 cm of the horizontal and vertical axes.

Electrophysiological data was sampled and stored at 30,000 Hz by the Cerebus data acquisition system (Blackrock Microsystems, Salt Lake City, UT, USA). Cerebus Central Suite software (Blackrock Microsystems) was used to visualize both neuronal and motor unit data, classify units online, and mark the time of action potentials.

### Data Analyses

Neural and EMG data was band-pass filtered (250Hz-5kHz) and sorted into individual units offline using the offline sorter provided with the Cerebus Central Suite (Blackrock Microsystems). Spike-timings obtained after offline sorting were down-sampled to 1 KHz to match the sampling rate of task parameters. Saccade onset and offset times were detected from the eye position data using a velocity threshold. All analyses, post spike sorting, were done using custom-made scripts written in MATLAB (MathWorks, Natick, MA, USA). The final dataset used in this study comprises units primarily showing presaccadic activity: 38 FEF movement neurons and 76 motor-units. To compare across the neuronal and EMG data, both types of data were displayed as continuous spike density functions (SDF) and were analyzed similarly. The spike density functions were calculated by convolving the averaged spike train with a filter that resembled an excitatory post-synaptic potential, having a combination of growth and decay exponential functions. The time constants of the rapid growth phase (τ_g_ = 1 ms) and the slower decay phase (τ_d_ = 20 ms) were matched to values obtained from excitatory synapses (Kim and Connors, 1993; Sayer et al., 1990).

#### Response field

Response field (RF) identification was done using the memory-guided saccade task. Three target locations with the highest activity were named as locations inside the response field and the three diametrically opposite locations were considered to be outside-RF locations. The first target in the FOLLOW task could appear at any of six inside response field and outside response field locations. The second target in step trials, however, could only appear at any of the three positions diametrically opposite to the location of the first target. This was done to maintain a wide separation between the two saccade targets and prevent averaging of the first and second saccades.

#### Visuo-motor index

We classified the neurons and motor units by identifying their discharge patterns in the memory-guided saccade task. We also computed a visuo-motor index (VMI) to quantify the ratio of target-related and saccade-related activities among the classified neuron (Murthy et al., 2007).

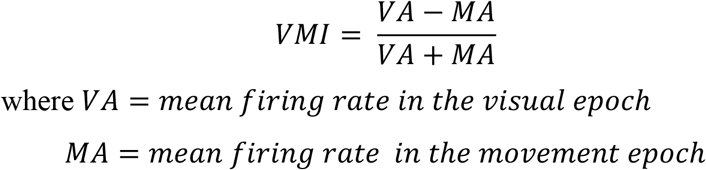

VMIs ranged from +1 to -1. Movement neurons had highly negative VMIs (<-0.33).

#### Measures of EMG activity dynamics

We analyzed the EMG data using an accumulator framework, looking at four main features: baseline, onset, rate of activity growth, threshold. The results were compared with similar parameters calculated in a prior study on FEF neurons (Basu et al., 2021). Each accumulator parameter was calculated separately for the first saccade plan (first saccade was made to the RF) and the second saccade plan (second saccade was made to the RF).

Differential activity (activity in step trials – activity in outside-RF no-step trials) was used to calculate accumulator parameters for both motor units and neurons.

- Baseline activity = Mean differential activity in the 100 ms period before onset of first target.
- Onset of saccade-related activity = First time-point at which differential activity exceeded two standard deviations (SD) of baseline activity, ultimately crossing 4SDs, and staying above 2SDs for 50 ms (second plan) and 30 ms (first plan).
  ∘ The onset of second saccade-related activity or the peripheral selection time (PST; shown in **Fig. 3**) was calculated similarly (Basu and Murthy, 2020). PST was defined as the time point when differential activity in trials where the second saccade went into RF exceeded 2 SDs above baseline activity and stayed above 2SDs for 45 ms and reached 4SDs within this time window.
- Threshold activation: Mean activity in the 10 to 20 ms period preceding saccade onset.
- Growth rate: Difference between threshold activation and activity at onset, divided by the time period from onset to threshold.

For visualizing population activity profiles, SDFs were normalized to the peak activity in the TSD 17 ms condition for each motor unit.

### Statistical tests

For a single group of data, a two-sided Wilcoxon signed-rank test was used. For comparison across multiple groups, the Kruskal-Wallis test was primarily used. All the results are presented as mean (± standard error of mean, SEM) and all tests are performed at a significance level of α = 0.05 unless otherwise mentioned.

## Supplementary Figures

**Figure S1.**
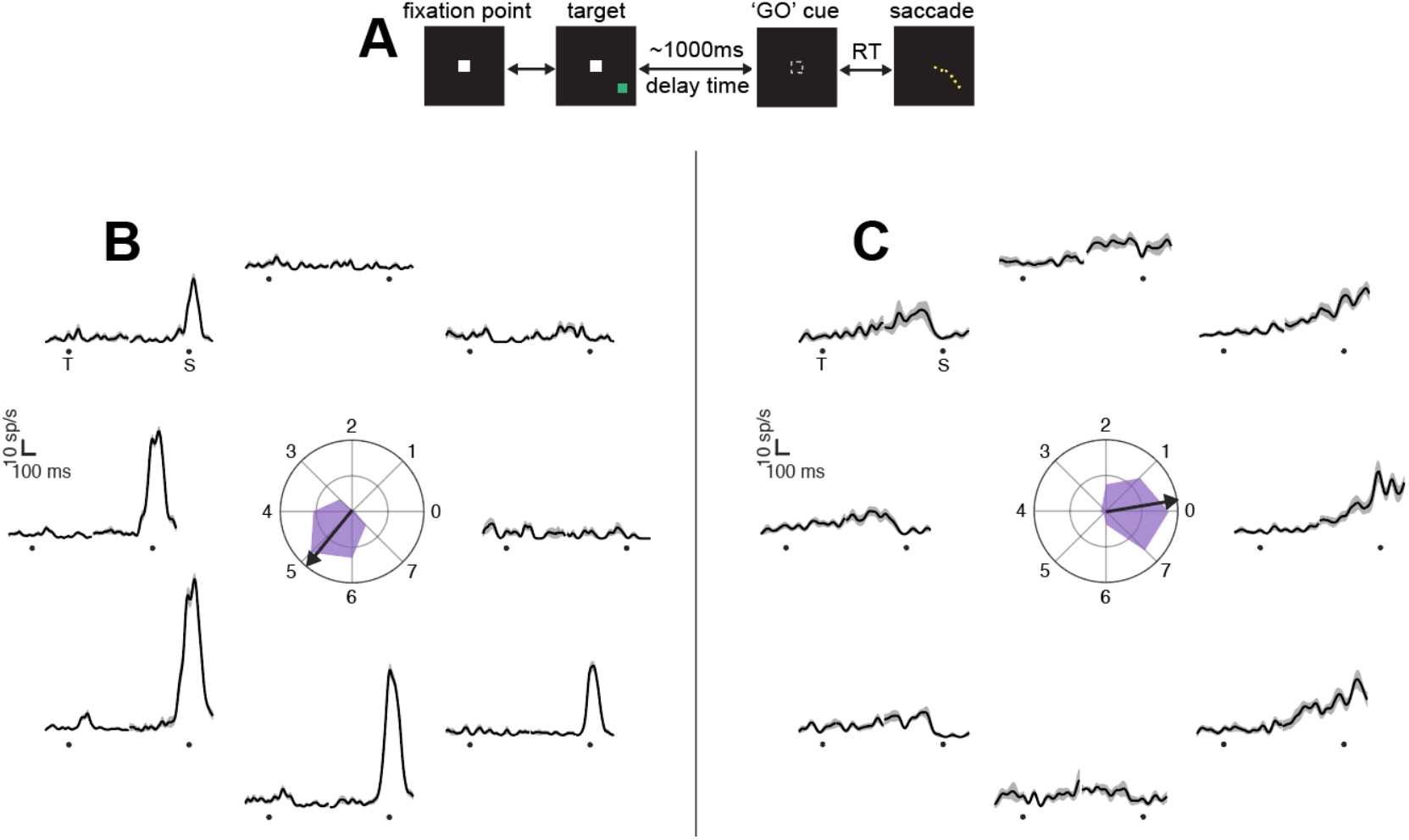
Memory guided saccade task and electrophysiology. **A**. Schematic of the memory-guided (MG) saccade task. **B**. Activity of a representative FEF movement neuron in the MG task for all eight target positions. **C**. Same as **B** but for a representative motor unit.

**Figure S2.**
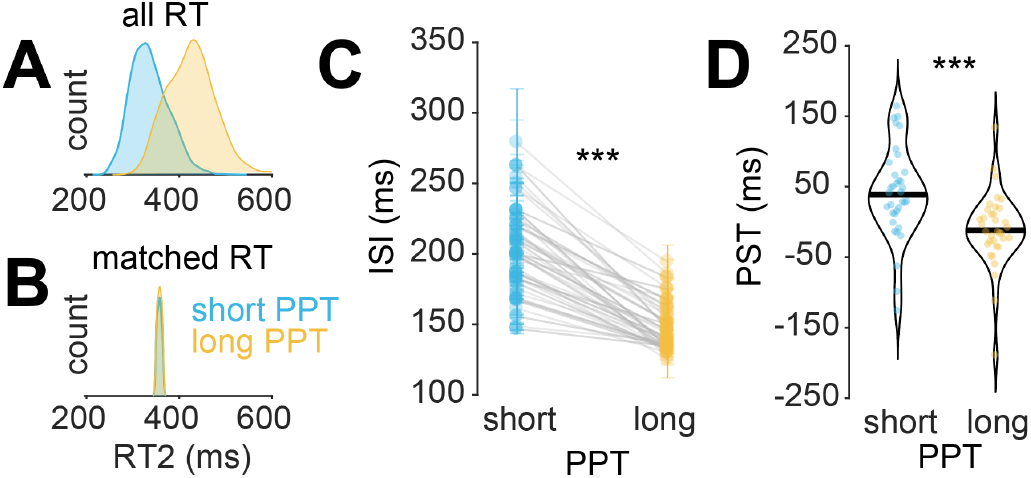
Activity of motor units show signatures of parallel programming in latency-matched trials. **A**. Distribution of second saccade reaction times for the short (*blue*) and long (*yellow*) PPT conditions. **B**. Reaction times that were matched between the short and long PPT conditions. **C**. ISIs vs PPT plot for reaction time matched trials. *** means *p* < 0.001 **D**. Peripheral selection times for short and long PPT conditions for latency-matched trials. *** means *p* < 0.001

**Figure S3.**
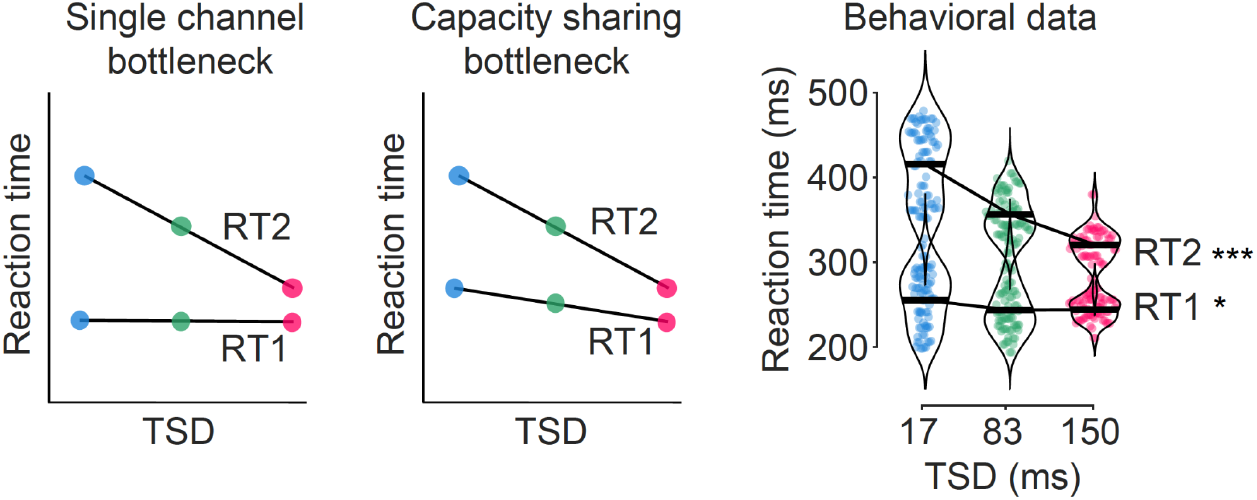
Reaction time vs target step delay. *Left panel*: Reaction time vs target step delay (TSD) predictions based on single-channel bottleneck model. *Middle panel*: Reaction time vs target step delay (TSD) predictions based on capacity-sharing bottleneck model. *Right panel*: Behavioral data for reaction time vs target step delay (TSD). Data shows trials in which the first (for RT1) or second (for RT2) saccade was into the response field. Both reaction times increased significantly with decrease in target step delay. *** means *p* < 0.001 and * means *p* < 0.05

**Figure S4.**
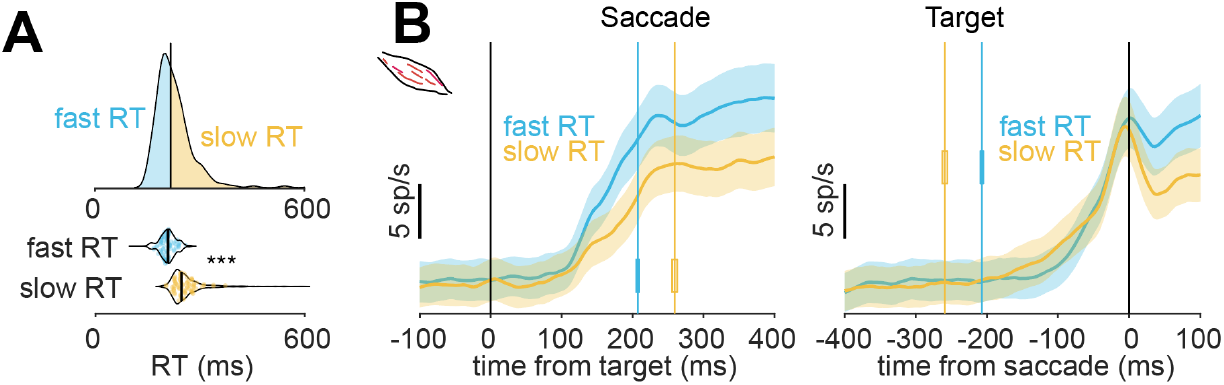
Activity of motor units correlated with reaction time in no-step trials. **A**. *Top*: Reaction time (RT) distribution for a representative session showing the fast and slow reaction times. *Bottom*: Fast and slow reaction time values for all recorded sessions. *** means *p* < 0.001 **B**. EMG activity in single saccade no-step trials ramped up faster for saccades with faster reaction time (*left*). The threshold activity did not change across the two conditions in the population (*right*). The solid line indicates the mean activity and the shaded area indicates mean ± SEM.

